# Inference of genomic spatial organization from a whole genome bisulfite sequencing sample

**DOI:** 10.1101/384578

**Authors:** Emanuele Raineri, François Serra, Renée Beekman, Beatriz García Torre, Roser Vilarrasa-Blasi, Iñaki Martin-Subero, Marc A. Martí-Renom, Ivo Gut, Simon Heath

**Affiliations:** CNAG-CRG, Centre for Genomic Regulation (CRG), Barcelona Institute of Science and Technology (BIST), Baldiri i Reixac 4, 08028 Barcelona, Spain.; Biomedical Epigenomics Group, Institut d’Investigacions Biomédiques August Pi i Sunyer (IDIBAPS), Barcelona, Spain.; ICREA, Pg. Lluís Companys 23, 08010, Barcelona, Spain; Universitat Pompeu Fabra (UPF), Plaça de la Mercè 10, 08002, Barcelona, Spain; Gene Regulation, Stem Cells and Cancer Program, Centre for Genomic Regulation (CRG), Dr. Aiguader 88, 08003, Barcelona, Spain

## Abstract

Common approaches to characterize the structure of the DNA in the nucleus, such as the different Chromosome Conformation Capture methods, have not currently been widely applied to different tissue types due to several practical difficulties including the requirement for intact cells to start the sample preparation. In contrast, techniques based on sodium bisulfite conversion of DNA to assay DNA methylation, have been widely applied to many different tissue types in a variety of organisms. Recent work has shown the possibility of inferring some aspects of the three dimensional DNA structure from DNA methylation data, raising the possibility of three dimensional DNA structure prediction using the large collection of already generated DNA methylation datasets. We propose a simple method to predict the values of the first eigenvector of the Hi-C matrix of a sample (and hence the positions of the A and B compartments) using only the GC content of the sequence and a single whole genome bisulfite sequencing (WGBS) experiment which yields information on the methylation levels and their variability along the genome. We train and test our model on 10 samples for which we have data from both bisulfite sequencing and chromosome conformation experiments and our most relevant finding is that the variability of DNA methylation along the sequence is often a better predictor than methylation itself. We then run a prediction on 206 DNA methylation profiles produced by the Blueprint project and use ChIP-Seq and RNA-Seq data to confirm that the forecasted eigenvector delineates correctly the physical chromatin compartments observed with the Hi-C experiment.

## 1 Introduction

There are a number of techniques available to ascertain the 3D conformation of the genome in the nucleus. Hi-C sequencing [1] uses the frequency by which fragments of DNA are cross-linked, ligated and successively sequenced together as an indication for their proximity; the output of this technique can be represented as a square interaction matrix. Knowledge of the first (i.e. associated to the largest eigenvalue) eigenvector of the corresponding covariance matrix allows one to reconstruct a rough approximation of the interactions [1], [2]. An example of such an eigenvector is given in Figure 1.

**Figure 1:**
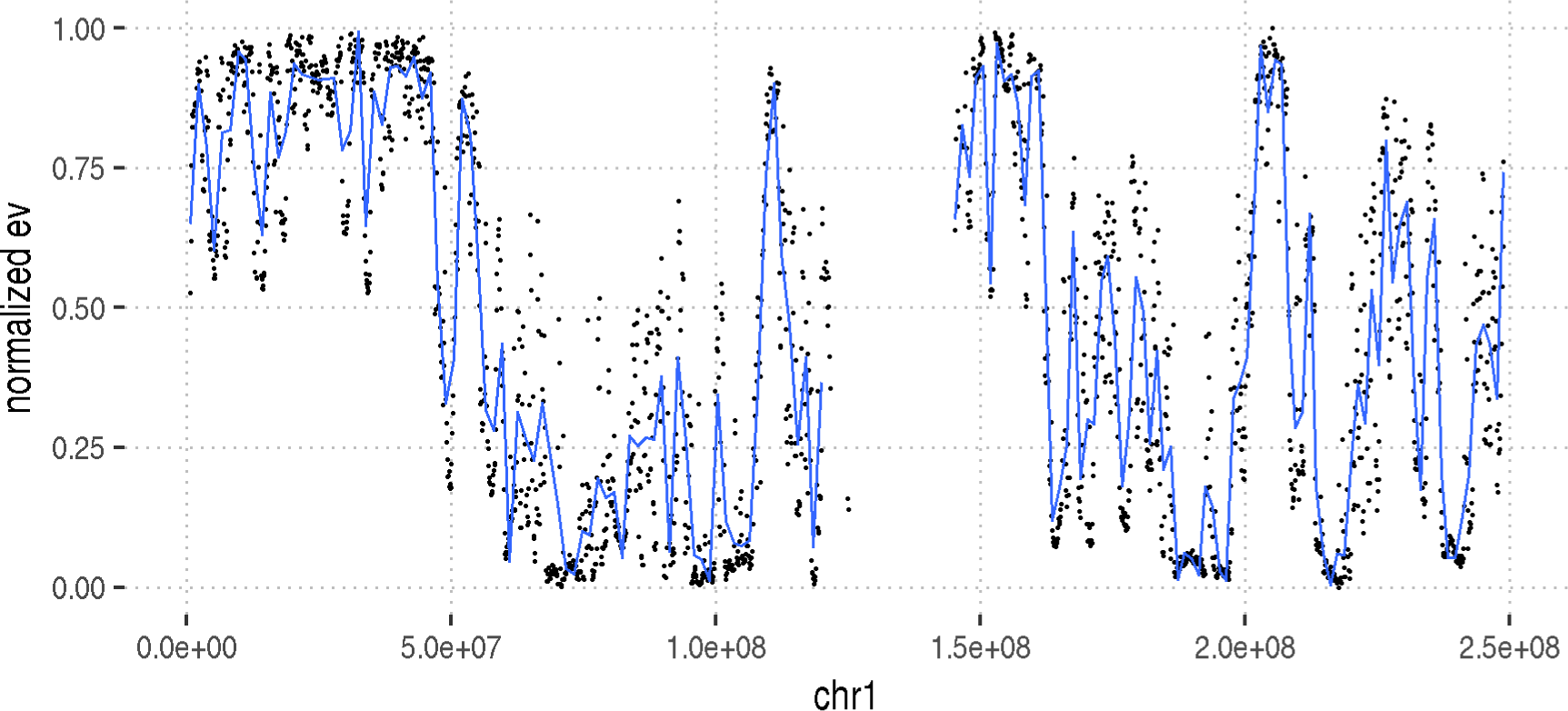
First eigenvector (black dots, with smoothing in blue) of the Hi-C matrix as measured on chr1 of a memory B cell at 100 kb resolution. There are two distinguishable low and high states.

The folding of the genome is associated to other epigenetic signals which can be exploited to predict it [3]; for example in [4] the authors use random forests on ChIP-Seq data to infer Hi-C matrices; Huang and others employ Bayesian additive regression trees for a similar purpose in [5]. It has also been shown that DNA methylation alone could be used to infer one of the most important expression of genome folding, A/B compartments [1]. This predictive power of DNA methylation was described for the first time by Fortin et al. [6] using hundreds of methylation 450k microarrays. Simmonds et.al. [7] also use methylation 450k microarrays to predict A/B compartments and to study differences between healthy and cancerous genomes. Jenkinson et al. [8] classify compartments from a single WGBS experiment by way of segmenting DNA methylation via an Ising model.

In this work, we also set out to use only the information provided by DNA methylation; our model differs from the work of [6] in that we use one WGBS sample only as opposed to tens or hundreds of microarray; and differs from [8] for a number of reasons: we use a simple model with negligible computational cost, we predict directly the eigenvector as opposed to the A/B compartments, we test our model on more cell types; finally, we distinguish the information brought by the methylation level, its variability along the genome, and the GC content of the genome itself. Our model is accurate enough that repressed and active regions can be determined correctly in many cases: for example the correlation between transcription and predicted eigenvector is around 0.6-0.7. The fact that we estimate directly the eigenvector (rather than only the compartments), might allow for the analysis of intermediate structures (to this respect see for example [9] where Rao, Huntley and others argue that there are in fact 6 types of subcompartments). In what follow we will also explore the applicability of this approach to 450k array and high resolution Hi-C data. Finally, we validate our predictions using RNA-Seq and Chip-Seq measurements.

## 2 Materials and Methods

### 2.1 Training

We train linear regressions with different covariates on the transformed first eigenvector of the Hi-C matrix and compare their performance when predicting. The dataset adopted for fitting models (for each chromosome and replicate), contains 10 Hi-C eigenvectors (2 from experiments performed described in [10] and processed through the TADbit pipeline [11] and 8 provided to us by the authors of [12]), and 10 matching WGBS experiments ran by the Blueprint consortium (see table 1) The eigenvectors are measured at a resolution of 100 kb; since the samples studied in [12] were originally mapped to hg19 we convert the coordinates to hg38 (for consistency with the other datasets we harness) using the liftOver tool [13], hence in some cases the resulting windows are not 100 kb long. Figure 2 shows how the eigenvectors (on chromosome 1) cluster according to cell type.

**Figure 2:**
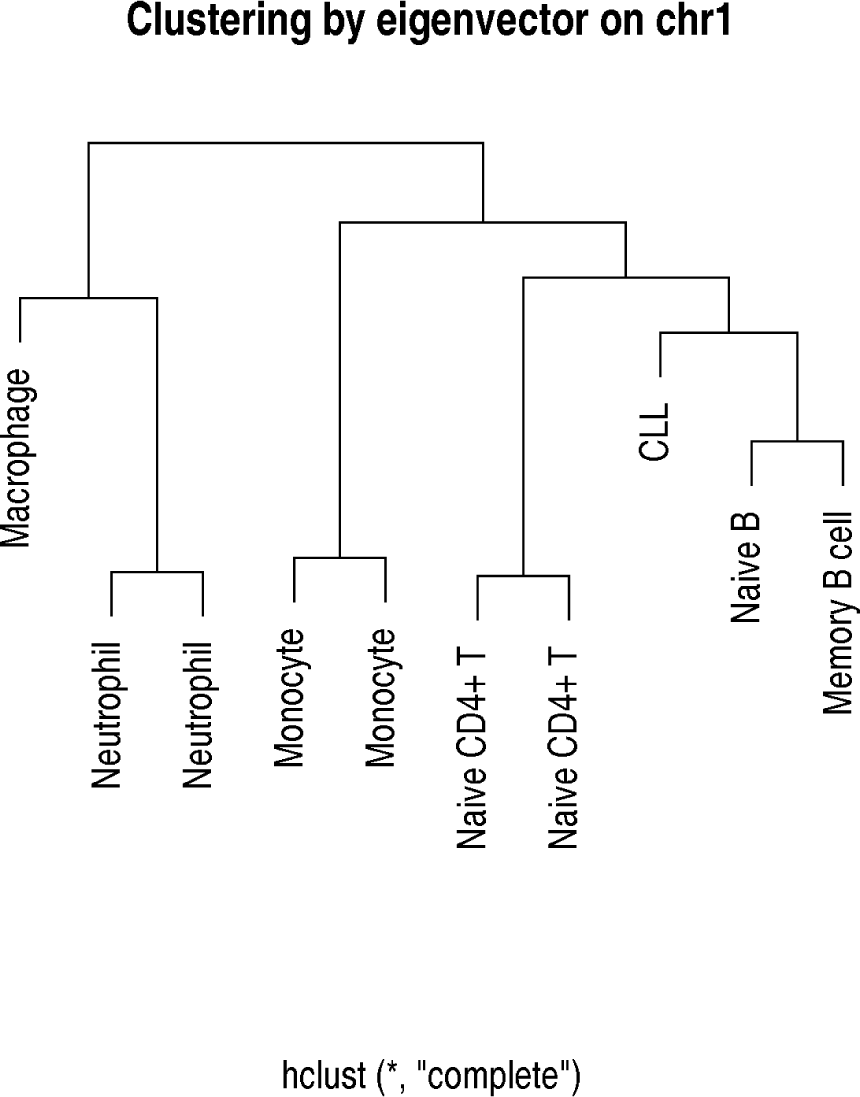
Hierarchical clustering of measured eigenvectors both taken from [12] and [10].

**Table 1:**
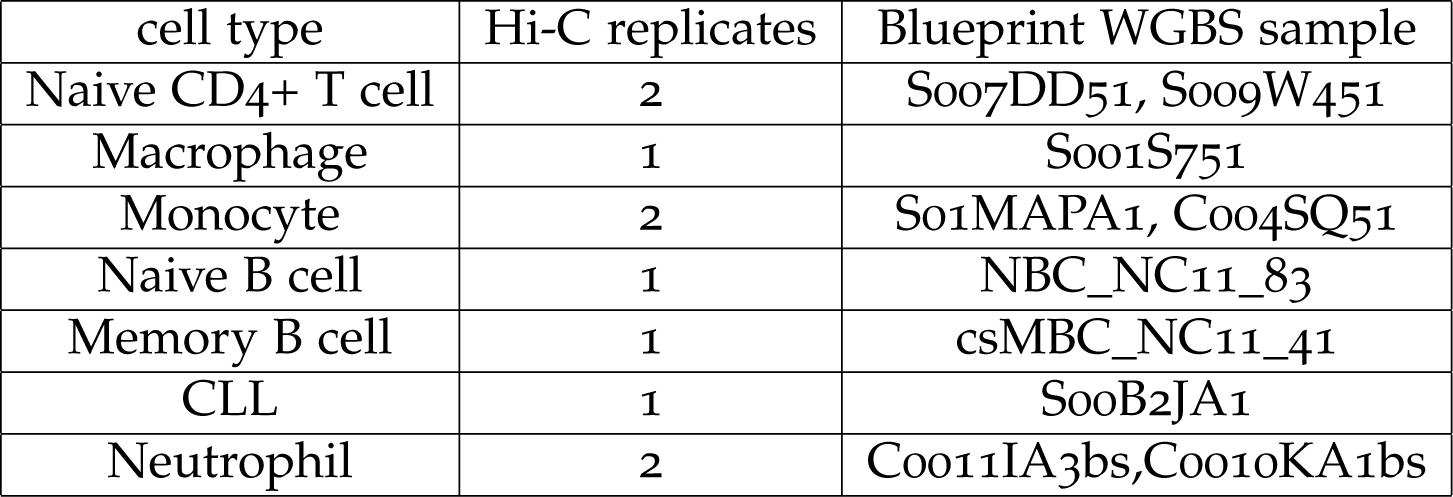
Samples used for training.

To prepare the data for the statistical analysis, we first divide the genome into adjacent (non overlapping) windows of 100 kb (the analysis was also repeated at 10 kb and 50 kb, see below). For each window we consider the median methylation level x_m_, the standard deviation of methylation x_sdev_, the number of C or G nucleotides per base pair x_GC_. We also initially looked at the fraction of CpG dinucleotides per base pair x_CpG_ but we find it does not provide any predictive power once the GC content is taken into account. The estimation of GC content can be inaccurate if the window contains many unidentified bases (N). For this reason windows with more than 10% of Ns are excluded. Also, one should remember that we are dealing with populations of cells: if we disregard allele-specific phenomena, a genomic position can only have methylation 0 or 1. The intermediate values we are discussing here are due to the presence in the sample of various subgroups of cells which are methylated or unmethylated at a given position. Figure 3 contains plots of each covariate against the others, and against the normalized training eigenvector for chr1 across all training samples. Since the sign of the eigenvectors is arbitrary we choose it so that it is positively correlated with GC content; we also standardize them by rescaling them to the interval [0, 1]. In what follows the eigenvector will be designated by Y and we will work with the transformed version 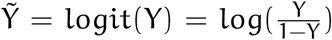 because it results in a better fit of the models below. We do not consider interactions between covariates ( i.e. the model is purely additive, it does not contain products of covariates) as they seem to have a weak effect; consequently there are 8 possible design matrices corresponding to the possible combinations of covariates in the equations. For example (if we let 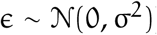) one possible choice corresponds to the equation 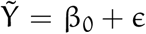 (no covariates, the intercept) whereas another could be 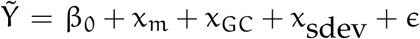 only (all covariates) and so on. We will not use mathematical notation when referring to the models in the rest of the paper and talk instead of the (for example) GC+meth+sdev regression. The regression with no covariates apart from the intercept will be used throughout the paper as a term of comparison; it is the choice which has the maximum prediction error, as it should be intuitively because it does not take into account any information about the genome. In fact, it corresponds to a constant predictor which always returns the average value of the training eigenvector at all positions. As such, it will provide context for the plots, as it will show how much we gain with the knowledge of various experimental measurements. As a measure for the performance of the regressions we will use the median of the absolute differences with respect to the true eigenvector. This is a robust measure, in the sense that a few outliers in one direction or another are unlikely to sway the median.

**Figure 3:**
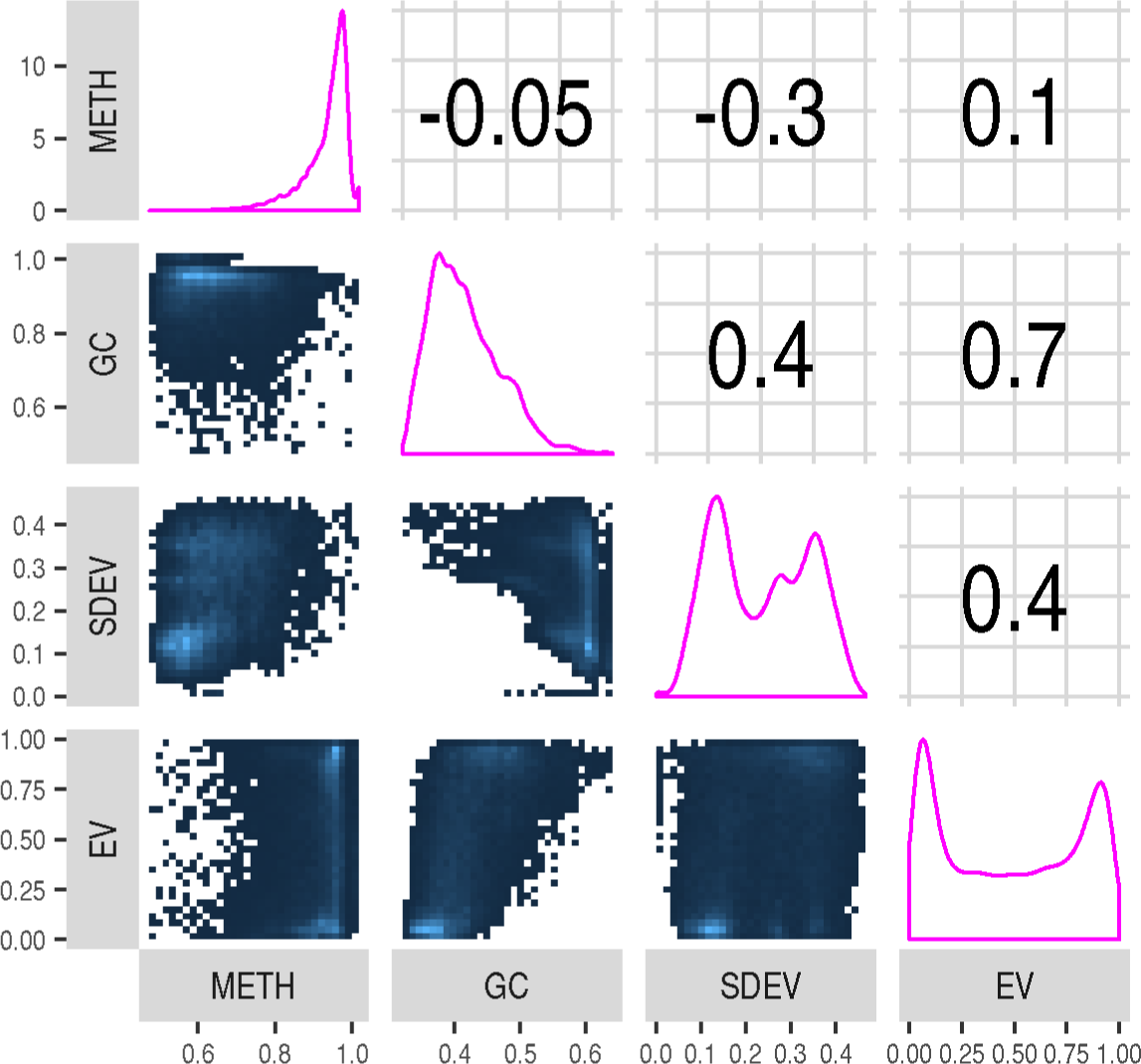
Covariates plotted against each other and the normalized eigenvector on chromosome 1 across all the training samples. The numbers in the boxes are correlation coefficients.

We also consider (for one cell type only: memory B cells) eigenvectors computed (by the TADbit pipeline) at 10 kb and 50 kb; we use those to train and cross-validate as usual.

It is also useful to evaluate this approach on data which are sparser than a WGBS sample. To this purpose, we considered microarray data: 5 class switched Memory B cell (described in [14]) and 2 multiple myeloma samples (described in [15]). In both cases we trained using the eigenvector measured in memory B cells.

### 2.2 Validation

To validate the models we infer the eigenvector in samples for which we have no Hi-C data. To this purpose we train the GC+meth+sdev regression (which is overall the best one: see the Results section) on each chromosome as before but pooling the data across all the cell types (instead of training a regression for each combination of chromosome and cell type). In this way we aim at using one model which is not related to any specific cell type. In using a linear regression we are implicitly assuming that the residuals will be homoskedastic and normally distributed. These assumptions are never met perfectly by any real data set, but deviating from them too strongly might make the interpretation of the results difficult. To depict the goodness of fit we prepared a Q-Q plot [16] i.e. a plot of the standardized residuals of the aggregated model vs a standard normal distribution (see figure 4) which shows broad agreement but also a sizeable number of outliers. Possibly adding a nonlinear part to the model would diminish the outliers, but we prefer to accept them and not do ad-hoc changes.

**Figure 4:**
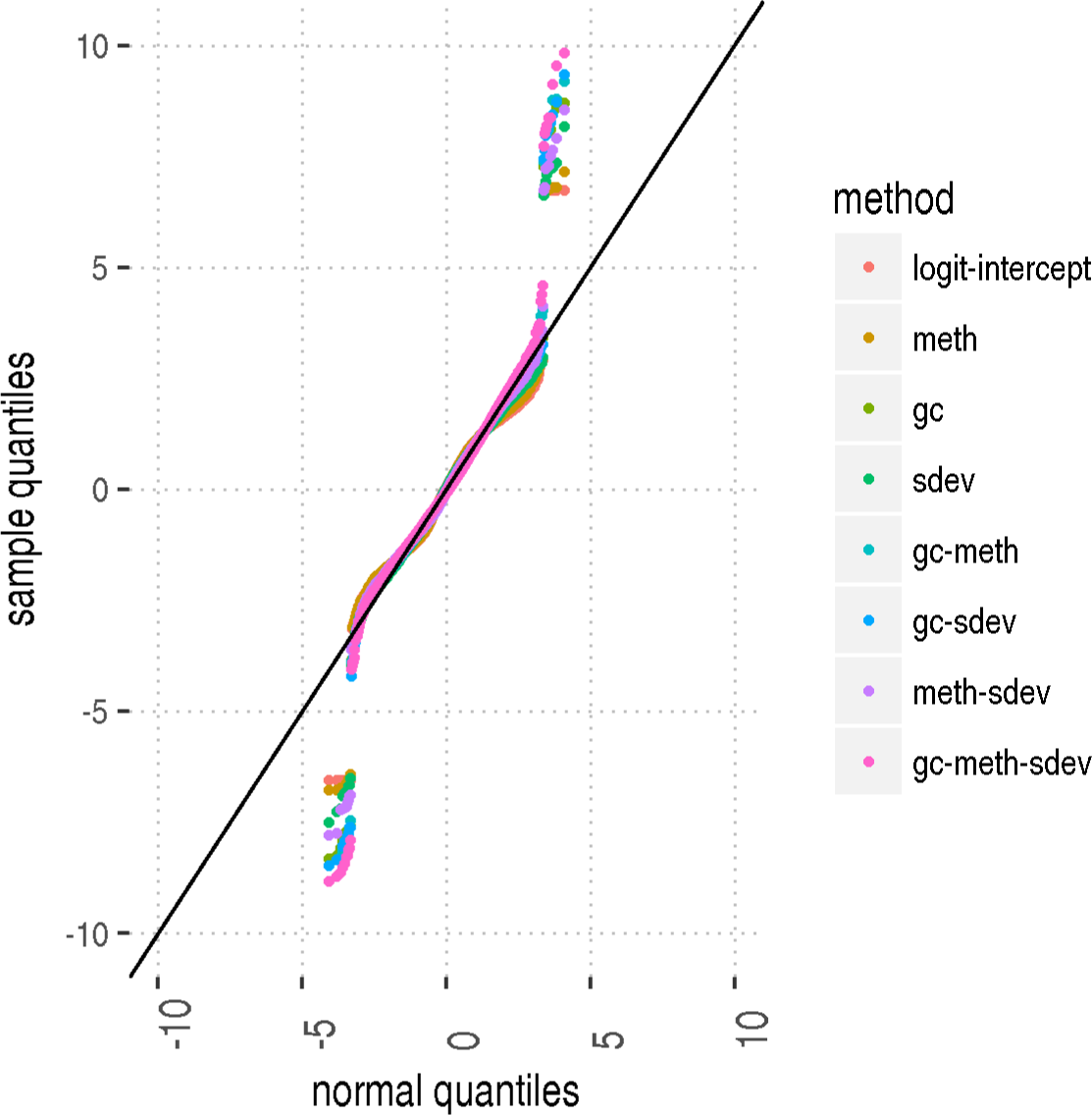
Quantiles of the standardized residuals (y axis) vs quantiles of a normal distribution (x axis), both expressed as Z scores. If the residuals were perfectly normal, the plot would coincide with the black diagonal line.

We also collected a set of measurements to test whether the predicted eigenvector is by and large reliable. Essentially we verify that activation signals (RNA-Seq, H3K27ac, H3K36me3, H3K27me3) and repression marks (H3K9me3) correlate as expected with the inferred eigenvector, where the prediction is made with the 8 models described above. We also check that the real eigenvectors used for training are correlated with such signals.

We collected relevant chromatin marks and RNA-Seq data as follows. First, for each WGBS sample we downloaded from the Blueprint repository (see http://dcc.blueprint-epigenome.eu/#/files) the coordinates of the chromatin peaks as measured in experiments matching as much as possible the WGBS one (for each methylation sample we consider the sex of the donor, cell type, tissue, cell line, disease status and the specific donor code; we select the chromatin experiment which matches the highest number of those). We count the number of peaks (with −log(pvalue) higher or equal to 3) falling in the windows mentioned above and correlate it with the predicted eigenvector. Furthermore, we fetch (with the same matching criteria) transcription quantification files which list TPM values for every gene. For each window we add the TPM of all the genes contained in it. When a gene is only partially contained in a single window we multiply its listed TPM value by the ratio of the length of the gene part which overlaps the window to the total gene length. We then compute the Spearman correlation between TPM value and predicted eigenvector.

## 3 Results

### 3.1 Training

At first, to characterize the quality of the models, we train on half the available data (by selecting at random half of the positions in each chromosome of each sample) and use the remaining half to compute the median absolute error (MAE) of the prediction. The boxplot in Figure 5 reflects the results of the exploratory analysis done using the 8 models above: GC content and standard deviation of genomic methylation are the best single predictors of the first eigenvector, followed by the median methylation value. Overall the best predictor was built using median methylation, GC content and sdev of methylation: it has a MAE which is significantly better than the GC predictor (difference in MAE 0.097 with pvalue (Wilcoxon) 2.1e-60). The observed error of the models varies depending on the cell type and chromosomes as shown in Figure 6.

**Figure 5:**
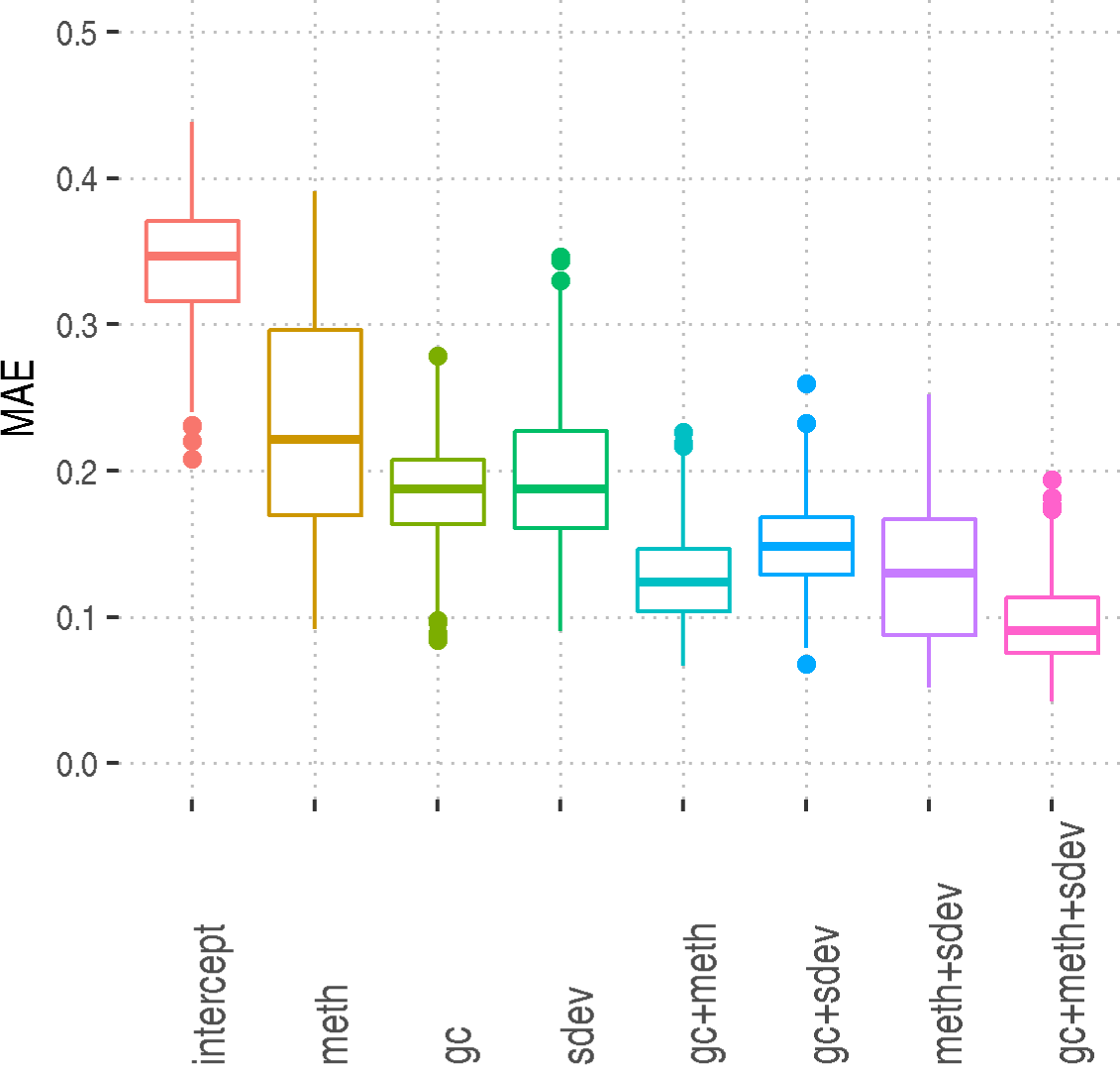
Median absolute error for some choices of covariates across chromosomes and cell types.

**Figure 6:**
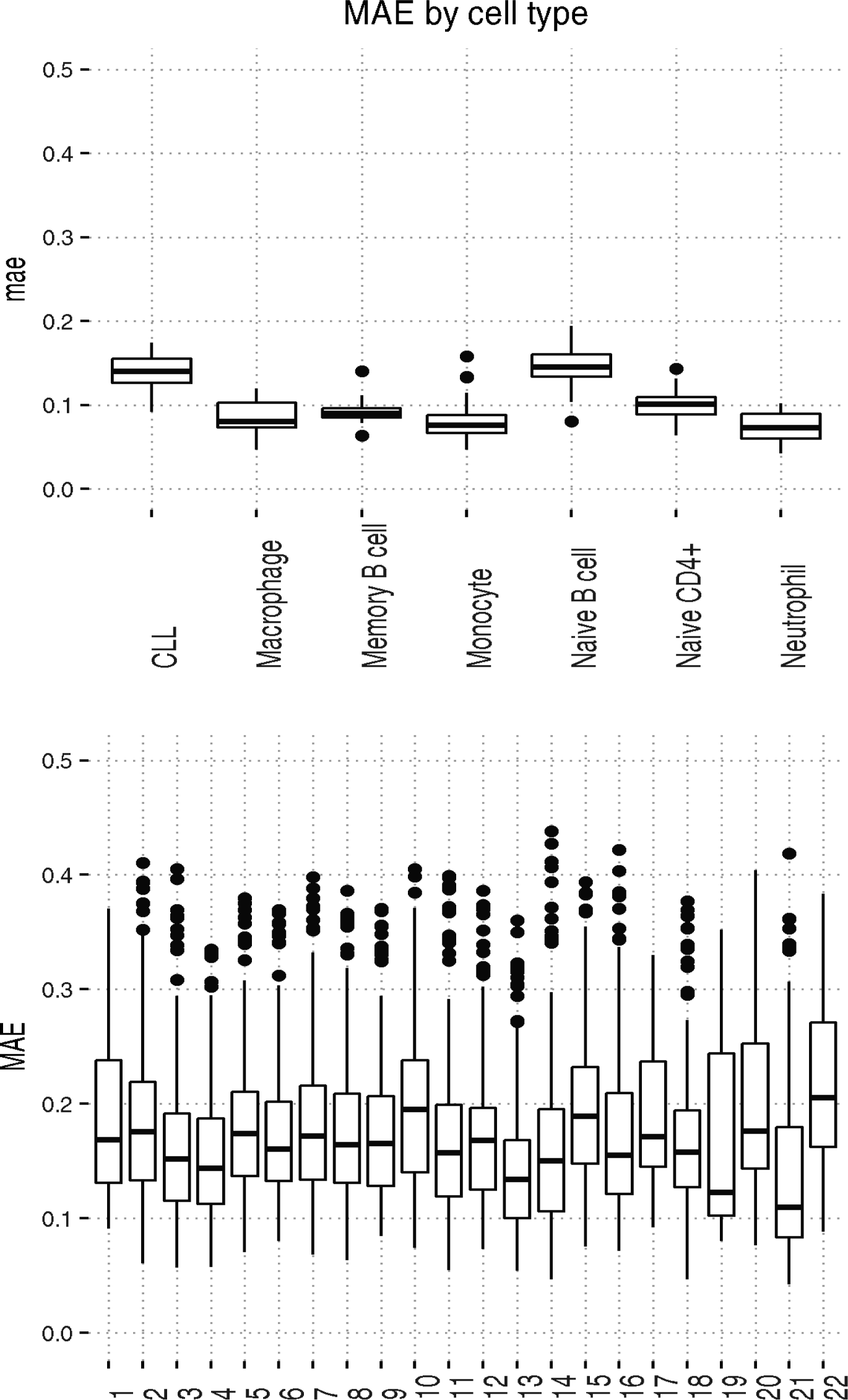
Top: Median absolute error of the gc-meth-sdev model for each cell type across all chromosomes. Bottom: Median absolute error for each chromosome across all models.

For what regards the differences in MAE due to cell type we noticed that Naive B cells seem more difficult to fit than Memory B cells (difference in median MAE is 0.056, p-value (Wilcoxon) 6.9e-07). This phenomenon might be caused by the fact that Naive B cells are heavily methylated (Figure 7), and therefore, the DNA methylation differences between euchromatin (A compartment) and heterochromatin (B compartment) are less marked than in memory B cells, which undergo a global demethylation in heterochromatic regions as explained in [14]. Correspondingly, figure 7 shows that methylation does not perform well as a predictor (on Naive B cells) while meth+gc+sdev does.

**Figure 7:**
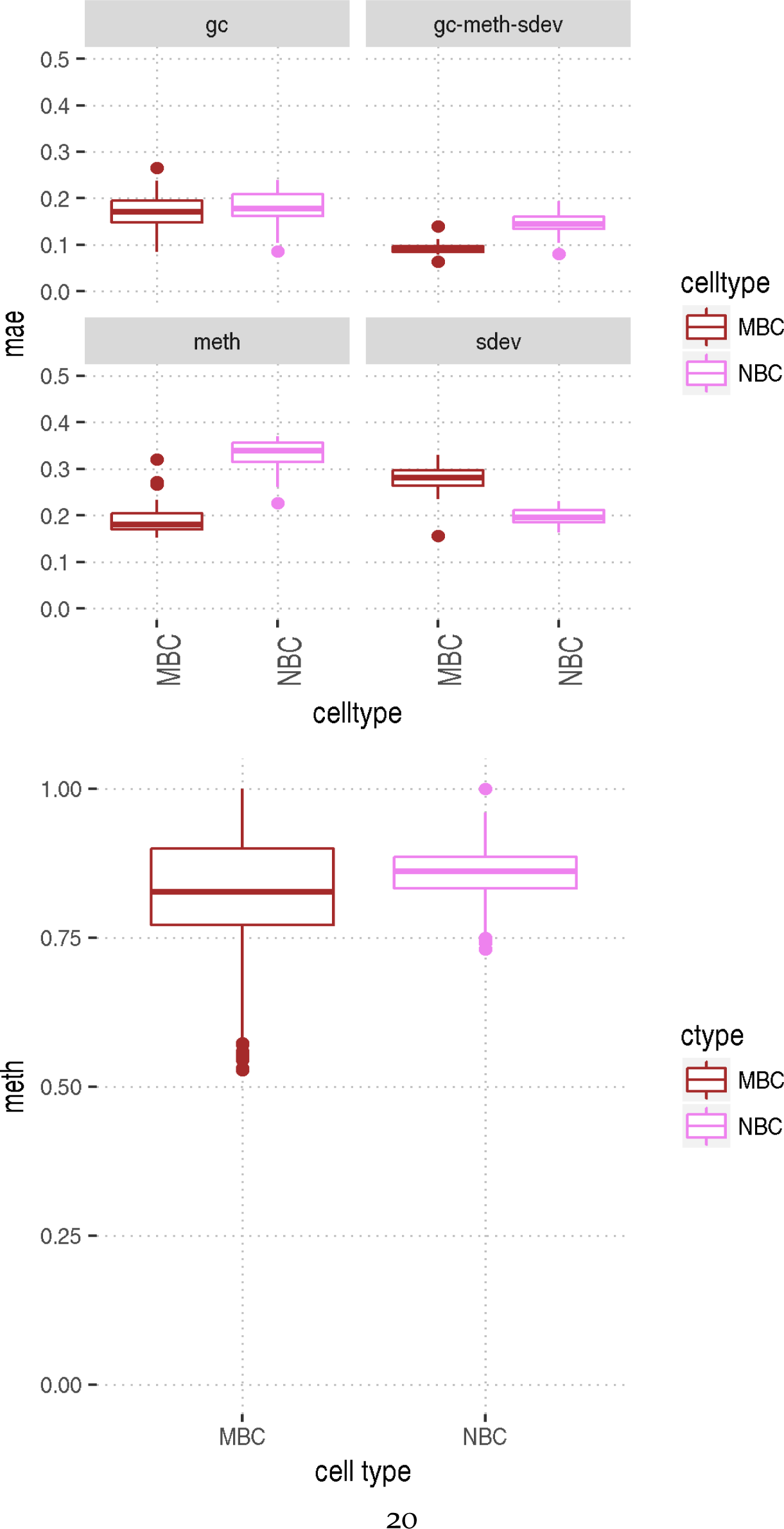
Top:in the Naive B cell sample the model including only methylation s a worse predictor than, for example, GC content. Bottom: comparison of methylation in windows of 100 kb for a Naive B cell (NBC_NC11_83, violet) and the memory B cell (csMBC_NC11_41, brown) used for training. In memory B cell, methylation is more variable.

When looking at Hi-C data at 10 kb and 50 kb we find that the best model at both resolutions was again meth+sdev+GC we also find that at 10 kb scale the prediction error is worse than the one obtained at 50 kb which is in turn worse than what we get at 100 kb (see Figure 8, which has the results chromosome by chromosome). It is difficult to say at this stage whether this is because our prediction method is particularly well suited for working at that resolution or because the eigenvectors computed at 10 kb and 50 kb are inherently less accurate than the 100 kb one. Also, we are examining one cell type only here. Nevertheless, the quality of the prediction does not degrade too much: when changing one order of magnitude in scale from 100 kb to 10 kb the median absolute error becomes roughly 1.5 times as big. For example for chromosome 10 it goes from ~0.1 to ~0.15.

**Figure 8:**
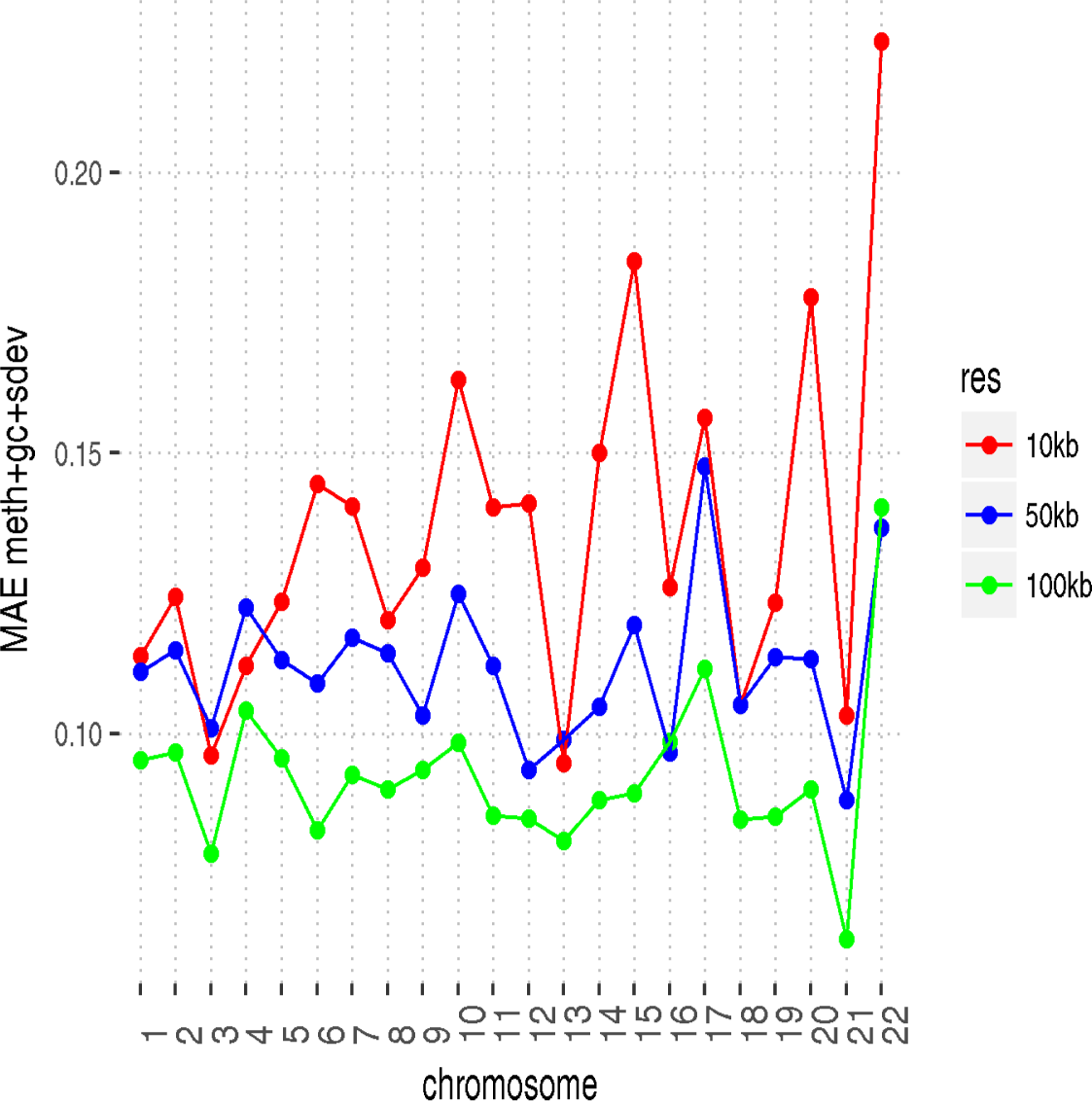
MAE for each chromosome at 10 kb,50 kb and 100 kb.

We cross validated the regressions on a microarray containing data on class switched memory B cells. The results (Figure 9) indicate that methylation and its standard deviation have some predictive power, but clearly less than the GC content. Also results in this case do not have a strong dependency on cell type; we always use a memory B cell for training but we show the results when predicting using both a memory B cell microarray and a multiple myeloma one: in the latter case errors are somewhat higher, but not by much. For example when using predictions made with gc+meth+sdev there’s a difference in MAE between memory B and multiple myeloma of −0.02 (pvalue Wilcox 0.13).

**Figure 9:**
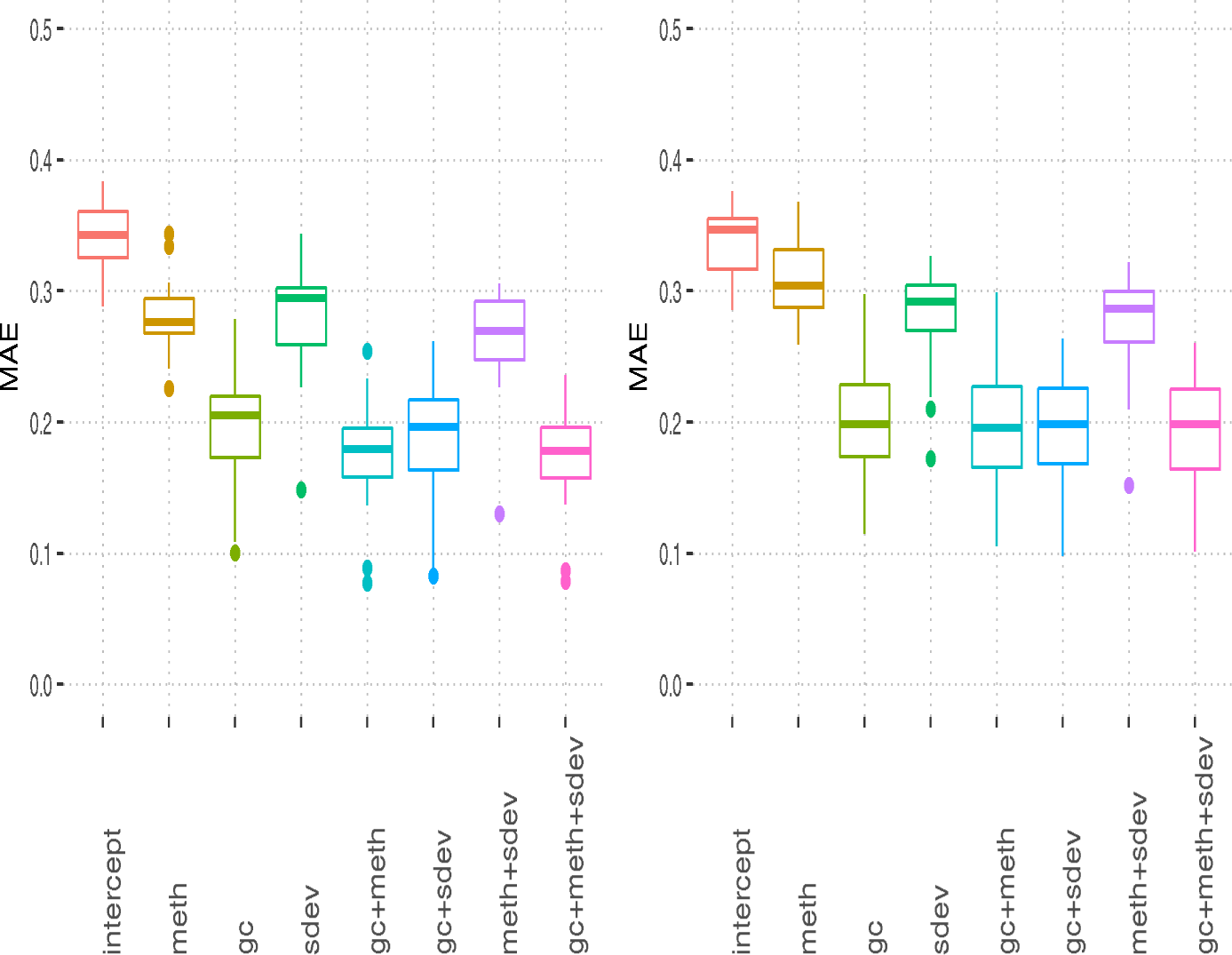
MAE when training on microarray methylation data. Left: MBC microarray, right multiple myeloma microarray.

### 3.2 Validation

We have been focusing so far on training and on measuring the cross-validation error. We will now test by predicting on new datasets (206 Blueprint WGBS samples, of which 10 used to learn the parameters of the regression) and using from now on a model which is averaged over the cell types as explained in the Methods section. First of all the pooled model can be verified on the Blueprint samples which we used for training: for example figure 10 shows the real and predicted eigenvector for one cell type for which we have both (Naive CD4+ T cell). For the rest of the samples, to make sure that our predictions are reasonable we check that they match active and repressed regions as defined by chromatin marks and RNA-Seq. We consider the 8 kinds of models described in the Methods section and we work under the null hypothesis that models which incorporate methylation and its standard deviation will produce forecasts as good or better than the ones obtained by looking at GC content only; we test this hypothesis by looking at the Spearman correlation between predicted eigenvectors and activation signals. This is the common theme of figures 11 and 12.

**Figure 10:**
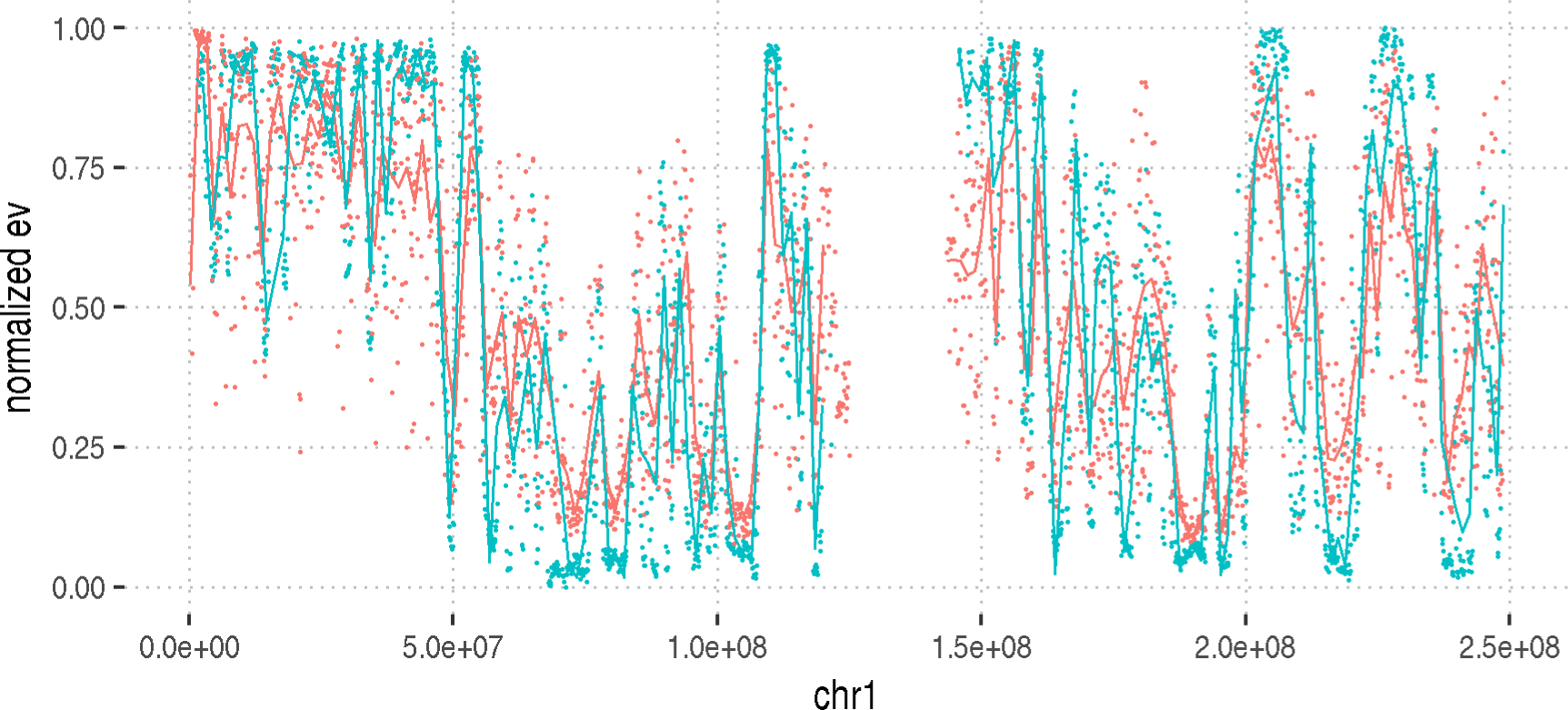
Comparison between predicted and measured eigenvector on chr1 of sample S007DD51 (Naive CD4+ T cell) at 100 kb resolution. Real eigenvector in blue, predicted eigenvector in red. In both cases we represent the values at each position and a smoothing line in the same colour.

**Figure 11:**
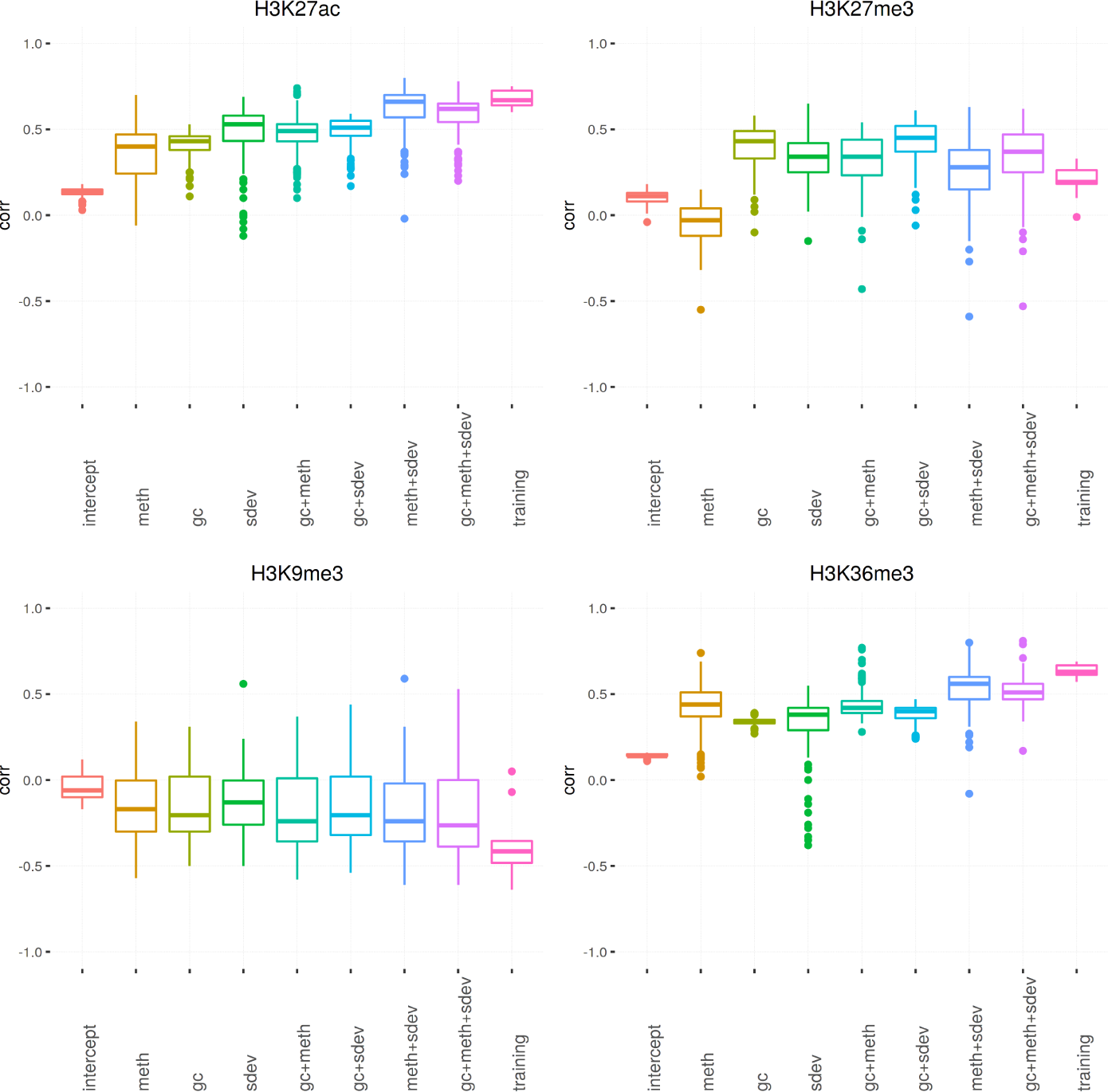
Correlation between predicted eigenvector and number of peaks for 4 chromatin marks and the 8 models considered and training eigenvectors. The box labeled as training corresponds to the 10 training samples, the other boxes to 206 validation samples.

**Figure 12:**
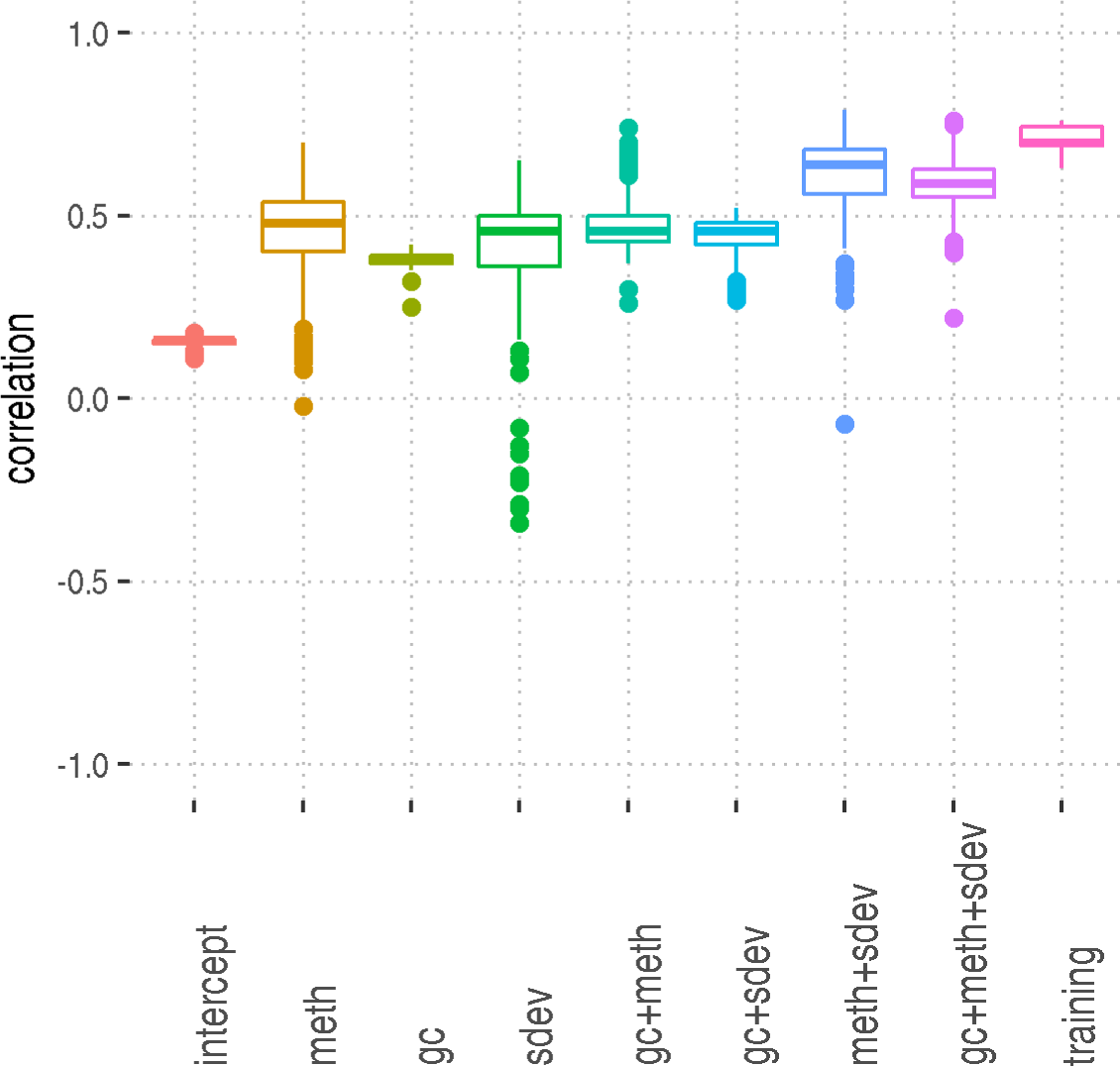
Correlation (Spearman) between the predicted eigenvector and the transcription levels per window, grouped according to prediction method and training eigenvectors. The box labeled as training corresponds to the 10 training samples, the other boxes to 206 validation samples.

In particular in figure 11 we first recapitulate the finding in [1] that both H3K27ac and H3K27me3 are positively correlated with the eigenvector, but H3K27ac more so than H3K27me3. We then investigate the behaviour of H3K9me3 as it is known to be associated with heterochromatin [17] and of H3K36me3, which is related to transcription (see figure 11). For what regards H3K27ac and H3K36me3, the correlations computed via the gc and the gc+meth+sdev are significantly different (for H3K27ac median MAE difference 0.19 with pvalue 4.6e-46 for H3K36me3 respectively 0.17 and 1.6e-66) the effect size is lower for H3K9me3 (median MAE difference −0.06 pvalue 0.00092 and H3K27me3 (median MAE difference −0.06, pvalue 0.0028).

We then look at the correlation between transcription and the predicted eigenvector, again contrasting different prediction methods. From figure 12 one can see that the gc+meth+sdev regression is close to the correlation obtained using real eigenvectors (difference of medians −0.11) and significantly different from the GC prediction (difference of medians 0.21, pvalue 1.9e-67). In figures 12 and 11 the boxplot for the real eigenvector is alway narrower than the others; this probably reflects the fact that we use only 10 samples for training (representative of 7 cell types) whereas we test against the Blueprint dataset which contains many more and more diverse cell types.

## 4 Discussion

We examined how to train and test models to infer a good approximation of the first eigenvector of the Hi-C matrix using only methylation and sequence information. The models need only one WGBS experiment to work, in contrast to many tens or hundreds of microarrays as in [6] and [7]; they are also simple to implement, unlike the more convoluted solution presented in [8]. Note that the form we chose for the regression does not imply causality: in fact, there are indications that methylation is established in already formed compartments [18]. WGBS experiments give two signals, the value of the methylation in a window, and its variability; they are both independently useful for predicting the 3D structure features of a region. Our figure 5 makes it clear that models which include information from bisulfite sequencing have a distinctly lower error than model incorporating only the GC content (difference in MAE 0.097); this is remarkable because GC content is the same for all cell types hence cell specific information might be encoded in the methylation and its variability. The fact that heterochromatic regions have higher standard deviation in DNA methylation (measured along the genome) is coherent with the observations that in late replicating regions (typically heterochromatic) the maintenance of methylation (i.e. the fact that it is copied during replication) is less efficient than in early replicating regions which might be due to reduced availability of substrate (or enzymes) at the end of the cell cycle [17]. Our predictions match active and repressed regions in the genome but different covariates contribute differently depending on the specific cell type, as we have explained in one specific example.

The applicability to 450k microarrays is probably limited by the sparsity of the data available there, as methylation adds little to the information provided by GC content (figure 9).

This strategy of using data that is relatively easy to produce to predict the results of more costly experiments has some practical appeal, but it also points the way to a simple quantification of the relation between 1D and 3D signal which is probably easy to extend and refine (for example by considering solo-WCGWs as described in [19]). This work could also help the interpretation of, or complement, RNA-seq experiments by pointing out which regions change their activity status across samples.

## 5 Funding

We acknowledge support of the Spanish Ministry of Science, Innovation and Universities (MICIU) to the EMBL partnership, of the Centro de Excelencia Severo Ochoa, of the CERCA Programme/Generalitat de Catalunya, of the Generalitat de Catalunya through Departament de Salut and Departament d’Empresa i Coneixement and of the co-financing by the Spanish Ministry of Science, Innovation and Universities (MICIU) with funds from the European Regional Development Fund (ERDF) corresponding to the 2014-2020 Smart Growth Operating Program. We also acknowledge support from the EU-FP7 project BLUEPRINT (282510).

FS and MR acknowledge the European Research Council under the 7th Framework Programme FP7/2007-2013 (ERC Synergy Grant 4D-Genome, grant agreement 609989) and the European Union’s Horizon 2020 research and innovation programme under grant agreement No 676556 (MuG).

## 6 Acknowledgments

We would like to thank Marta Kulis for sharing with us the microarray data used in this paper.

